# Chromatin regulates genome-wide transcription factor binding affinities

**DOI:** 10.1101/2022.04.04.486948

**Authors:** Hannah K. Neikes, Rik G.H. Lindeboom, Cathrin Gräwe, Lieke A. Lamers, Marijke P. Baltissen, Pascal W.T.C. Jansen, Simon J. van Heeringen, Colin Logie, Sarah A. Teichmann, Michiel Vermeulen

## Abstract

Transcription factor binding across the genome is regulated by DNA sequence and chromatin features. However, it is not yet possible to quantify the impact of chromatin context on genome-wide transcription factor binding affinities. Here we report the establishment of a method to determine genome-wide absolute apparent binding affinities of transcription factors to native, chromatinized DNA. Our experiments revealed that DNA accessibility is the main determinant of transcription factor binding in the genome, which largely restricts nanomolar affinity binding of YY1, SP1 and MYC/MAX to promoters, while FOXA1 also interacts with non-promoter elements with high affinity. Furthermore, whereas consensus DNA binding motifs for transcription factors are important to establish very high-affinity binding sites, these motifs are not always strictly required to generate nanomolar affinity interactions in the genome. Finally, we uncovered transcription factor concentration dependent binding to specific gene classes, suggesting transcription factor concentration dependent effects on gene expression and cell fate. Importantly, our method adds a quantitative dimension to transcription factor biology which enables stratification of genomic targets based on transcription factor concentration and prediction of transcription factor binding sites under non-physiological conditions, such as disease associated overexpression of (onco)genes.

## Introduction

Gene expression is regulated by a complex interplay between DNA sequence, chromatin structure and transcription factor binding^1–3^. A plethora of methods to characterize cellular epigenomes have been developed in recent years^4–6^. Furthermore, various methods to determine how transcription factors recognize DNA sequence motifs have been described^7,8^. DNA sequence, chromatin context, co-factors and DNA compaction are thought to have an important regulatory role on transcription factor binding and gene regulation^9,10^, while it is proposed that binding of different transcription factors is not all regulated in the same way^11^. Importantly, genome chromatization is predicted to regulate up to 98% of all transcription factor binding events in human cells^12^ and is, in addition to DNA sequence, the largest determinant of transcription factor binding^13^.

Observing where proteins bind in the genome does not explain *when* they bind certain loci. To biochemically understand transcription factor binding, the binding specificity and the affinity of a transcription factor^14^ for each DNA sequence must be determined. Binding specificity helps to predict which genomic sites are potential transcription factor binding sites, while the affinity determines for each genomic site at what transcription factor concentration it will be bound. In recent years, several *in vitro* techniques have been developed to determine global protein-DNA interaction specificities^8^, affinities^15^, or a combination of both^16^. Furthermore, many algorithms have been developed to predict transcription factor binding *in silico*, based on DNA accessibility, DNA sequence or gene expression data^17^. However, no existing technique is capable of quantifying absolute transcription factor binding affinities in the chromatinized genome (reviewed in^18^).

Here we present a method to determine Binding Affinities to Native Chromatin by sequencing (BANC-seq), in which native chromatinized DNA is used to determine genome-wide absolute transcription factor binding affinities. To this end, transcription factor concentration dependent binding to regulatory elements is quantified by either ChIP-seq or CUT&RUN^5^ to determine genome-wide binding affinities in a native chromatin context. BANC-seq enabled quantification of thousands of nanomolar apparent binding affinities for multiple transcription factors in multiple cell types, thereby allowing us to investigate the role of chromatin context and DNA sequence in regulating absolute genome-wide transcription factor binding affinities. BANC-seq adds a quantitative dimension to transcription factor biology, which confirms that chromatin context is the major determinant of transcription factor binding affinity^13^. Our data reveal that pre-existing, permissive chromatin architecture is a pre-requisite for high and low affinity transcription factor binding to occur. In addition, our results indicate that, by and large, the presence or absence of canonical transcription factor binding motifs differentiate high and low affinity transcription factor binding sites, respectively. Importantly, we show that chromatin context is interpreted differently by the pioneering transcription factor FOXA1 compared to YY1, SP1 and MYC. Finally, our data reveal that specific gene classes are bound at different transcription factor concentrations, suggesting concentration dependent effects on transcription and cell fate. These findings underscore the importance of incorporating binding affinities when investigating gene regulatory networks, and we therefore anticipate that BANC-seq will be an important tool for the gene expression, chromatin and quantitative systems biology community.

## Results

To determine genome-wide transcription factor-DNA binding affinities while retaining *in vivo* chromatin context, we established the BANC-seq workflow, in which intact chromatin is incubated with a titration-series of a purified epitope-tagged transcription factor (**Fig. 1a**). Next, the genomic binding sites at each transcription factor concentration are determined by either ChIP-seq or CUT&RUN, using an antibody against the tag-epitope. The nuclear isolation procedure is based on protocols for genome-wide DNA accessibility profiling, a well-established technique known to retain *in vivo* nucleosome positioning and transcription factor binding^4^. To investigate whether the BANC-seq workflow results in a loss of chromatin and co-factors for transcription factor binding, we used mass spectrometry to quantify the proteome before and after nuclear isolation. Importantly, while cytoplasmic proteins were lost, we observed no decrease in the abundance of nuclear transcription factors after nuclear isolation (**Extended Data Fig. 1a**).

**Figure 1.**
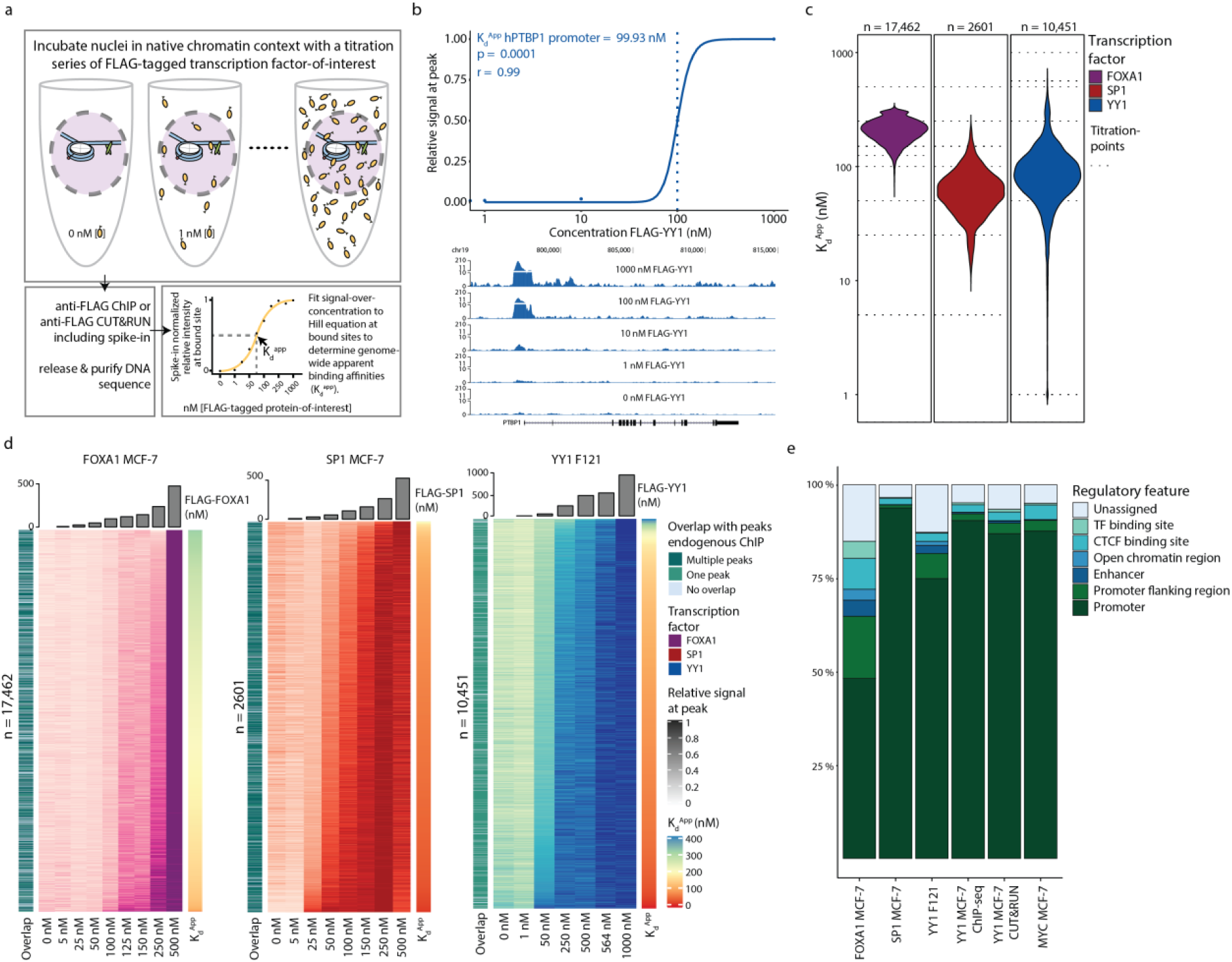
BANC-seq enables determination of genome-wide apparent binding affinities to native chromatin. (**a**) Overview of the experimental procedure. Isolated nuclei are incubated with a titration series of purified FLAG-tagged transcription factor, after which binding is quantified by anti-FLAG ChIP-seq or CUT&RUN, both including heterologous spike-in DNA. Spike-in normalised sequencing reads at each binding site and transcription factor-titration point are fitted to a Hill-curve to determine apparent binding affinities (K_d_^Apps^). (**b**) Spike-in normalised sequencing reads per titration point of FLAG-YY1 at the human PTBP1 promoter, relative to the highest signal (top; dotted line indicating the K_d_^App^), and visualized in the genome browser (bottom). (**c**) Distribution of K_d_^Apps^ of FOXA1 in MCF-7, SP1 in MCF-7 and YY1 in mESC cells F121. Dotted lines indicate the tested transcription factor concentrations per experiment. (**d**) Heatmap representing spike-in normalised sequencing reads relative to the highest signal for the same experiments as in (c). Each row represents one transcription factor binding site. The overlap of each binding site with peaks from endogenous ChIP-seq experiments of the same transcription factor is shown on the left of each heatmap, while K_d_^Apps^ are shown in the right column. (**e**) Overlap of identified binding sites with known regulatory features in the Ensembl Regulatory Build for different BANC-seq experiments.

Next, we set out to benchmark BANC-seq by adding 1, 10, 100 and 1000 nM FLAG-tagged YY1 to freshly isolated nuclei, followed by nuclear permeabilization, ten-minute incubation at 37°C and crosslinking. YY1 is a zinc finger transcription factor that binds enhancers and promoters to regulate gene expression and enhancer-promoter interactions^19^. Given the fact that the core sequence motif for specific YY1 binding is short, the amount of theoretical binding sites in the human genome is several orders-of-magnitude larger than actual binding sites observed by ChIP-seq. Therefore, we expected genome-wide YY1 binding affinities to be highly dependent on chromatin context. Sequencing revealed increased binding of YY1 at a known YY1 binding site with increasing transcription factor concentration (**Fig. 1b;** qPCR validation in **Extended Data Fig. 1b**). To quantify binding affinities, we used spike-in normalized ChIP-seq signal at the tested transcription factor concentrations to infer the apparent dissociation constant (K_d_^App^) at each identified binding site. To this end, we fitted parameters of a Hill-like curve to the observed signal over known transcription factor concentration (**Fig. 1a**; see *Methods*) and were able to determine high confidence apparent binding site affinities ranging from 36 to 156 nM for 5,372 genomic loci. Previously reported *in vitro* binding assays to determine affinities between YY1 and DNA have reported K_d_ values ranging from 3 to 4000 nM^20–24^, although most K_ds_ were found to be in the order of 100 nM, which is in good agreement with our measurements, providing a benchmark to profile additional transcription factors.

Encouraged by this pilot experiment, we performed BANC-seq with the pioneering transcription factor FOXA1, and transcription factors YY1 and SP1, and transcription factor heterodimer MYC/MAX in human breast cancer cells (MCF-7) and mouse embryonic stem cells. As ChIP-seq requires large amounts of cells as input material, we also adapted the protocol to be compatible with CUT&RUN to reduce the required number of input cells. These experiments revealed between 2,601 and 17,462 quantified binding affinities for per experiment (with a total of 48,220 quantified sites across the human and mouse genome), which span the entire nanomolar affinity range, equating to transcription factor expression of 100 to up to 100,000 molecules per nucleus (depending on the nuclear volume, see *Discussion*, **Fig. 1c-d**). The range of observed K_d_^Apps^ appeared to be dependent on the transcription factors we studied as well as and cell type (**Fig. 1c, Extended Data Fig. 2a-b**). However, using different methods (ChIP-seq and CUT&RUN) did not drastically influence the measured K_d_^Apps^, as one would expect (**Extended Data Fig. 2a**). Reassuringly, 83.6±7.6 % (mean ± SD) of the identified binding sites of exogenously added FLAG-tagged transcription factors overlapped with binding sites previously identified by endogenous ChIP-seq experiments for the tested transcription factors in the respective organism (**Fig. 1d, Extended Data Fig. 2b**), and 92.1±4.8% (mean ± SD) of the binding sites overlapped with annotated regulatory elements (**Fig. 1e**). This indicates that the exogenously added FLAG-tagged transcription factors exhibit physiological genomic binding in BANC-seq experiments, enabling faithful determination of genome-wide binding kinetics to regulatory elements by BANC-seq.

### Chromatin context impacts binding affinities

Next, we investigated the relationship between transcription factor binding affinities and chromatin context. We observed that almost all genomic sites for which we could determine high confidence transcription factor binding affinities are enriched for active histone marks, while being devoid of repressive and heterochromatin marks (**Fig. 2a**), illustrating that accessible chromatin is a prerequisite for transcription factor binding to occur. Nanomolar binding of SP1, YY1 and MYC/MAX appears to be almost exclusively restricted to promoter regions (**Fig. 1e**, mean ± SD = 86,7±0.71; and illustrated by the close proximity of peaks to transcription start sites depicted in **Extended Data Fig. 2c**). Strikingly, high affinity binding of these transcription factors is not often observed at enhancers, suggesting that chromatin is organized in such a way to facilitate binding of these transcription factors at promoters at low concentrations. Interestingly, low affinity binding sites for SP1, YY1 and MYC/MAX overlap more frequently compared to high affinity binding sites, suggesting that low affinity binding is largely controlled by DNA accessibility (**Extended Data Fig. 2d**). In contrast, nanomolar binding affinities for FOXA1 are not restricted to promoters (**Fig. 2a**): more than half of its quantified binding affinities are detected outside promoter regions, and identified binding sites are more distal to transcription start sites compared to those for the other tested transcription factors (**Extended Data Fig. 2c**). This observation may be explained by the fact that the pioneering factor capabilities of FOXA1^11,25^ make it less dependent on pre-existing chromatin architecture and DNA accessibility.

**Figure 2.**
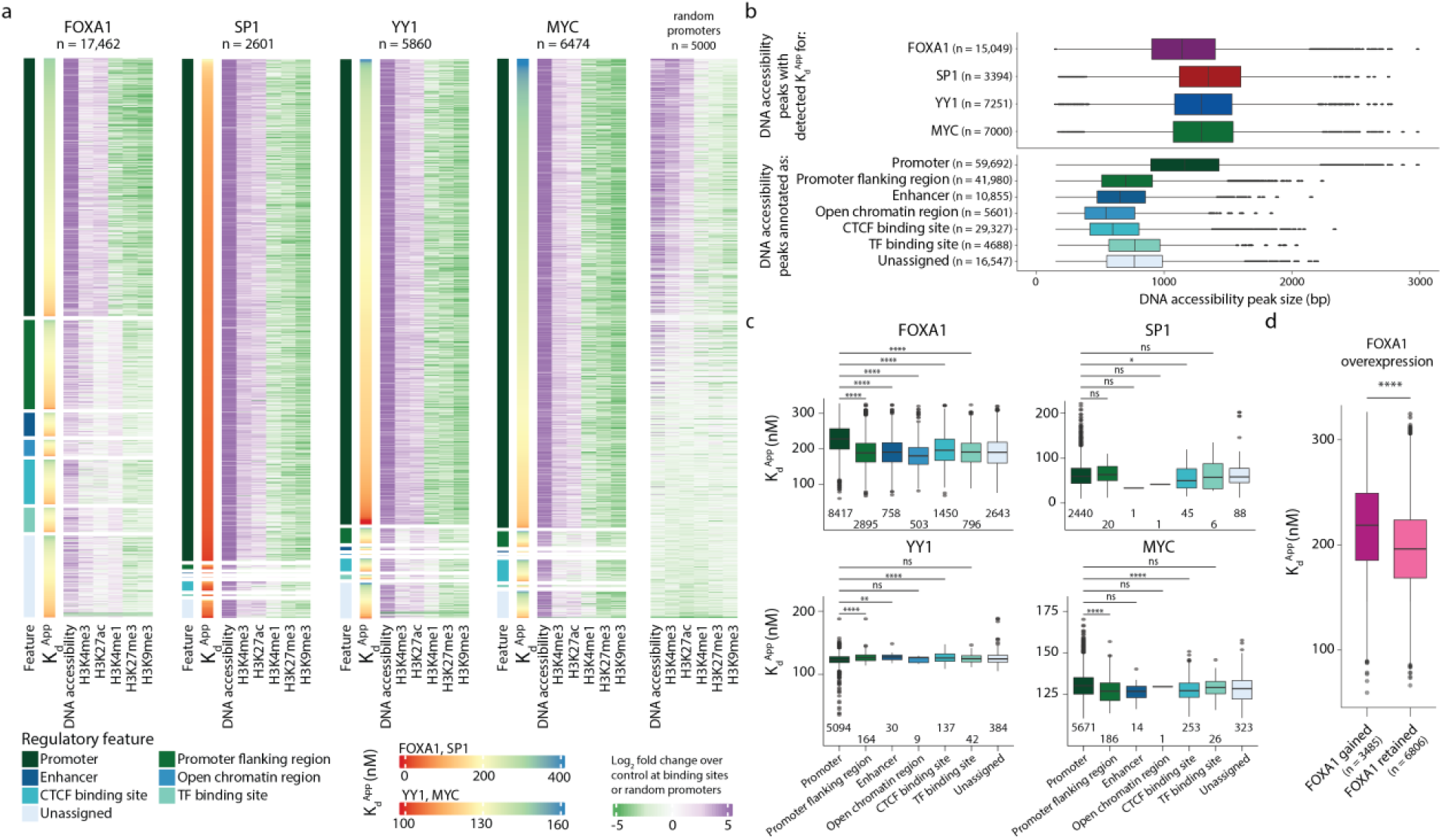
Chromatin context regulates transcription factor binding affinities. (**a**) Heatmap showing matched epigenome dynamics at binding sites with high-confidence K_d_^Apps^ fitted for FOXA1, SP1, YY1 and MYC in MCF-7, or 5,000 random promoters. Signals of ChIP-seq and DNA accessibility (by ATAC-seq) tracks for MCF-7 are shown as log_2_ fold change over the mean signal in five matched control tracks. Sites are ranked by assigned regulatory feature (first column per heatmap) and apparent binding affinity (second column heatmap). (**b**) Boxplots showing the size of DNA accessibility peaks at sites with high-confidence K_d_^Apps^ fitted per transcription factor (top), or per assigned regulatory feature (bottom). (**c**) Boxplots showing K_d_^Apps^ per assigned regulatory feature for the same transcription factors as in (a). Numbers at the bottom of each plot represent the number of sites in each group. (**d**) Boxplots showing K_d_^Apps^ for FOXA1 at sites that gain or retain FOXA1 binding after FOXA1 overexpression in MCF-7. **\*** are used to indicate significance according to a two-sided Wilcoxon test (ns: p > 0.05, *: p <= 0.05, **: p <= 0.01, ****: p <= 0.0001). Box plots were drawn with the center line as the median, the hinges as the first and third quartiles, and with the whiskers extending to the lowest and highest values that were within 1.5 × interquartile range.

To further investigate the correlation between DNA accessibility and high affinity transcription factor binding, we investigated whether promoters are more accessible in general compared to enhancer regions. Indeed, we observed that hyper-accessible regions in promoters are nearly twice as large compared to other types of regulatory elements (**Fig. 2b**), indicating that SP1, YY1 and MYC preferentially interact with larger accessible DNA elements. In contrast, FOXA1 bound sites are generally characterized by narrower and reduced accessibility (**Fig. 2b, Extended Data Fig. 3a**), possibly reflecting its ability to open condensed chromatin^26^. Here, FOXA1 was able to bind both promoters and enhancers with reduced accessibility, in particular at high affinity binding sites (**Extended Data Fig. 3a**), indicating that regulation of FOXA1 binding in the genome is fundamentally different compared to binding of SP1, YY1 and MYC.

In agreement with the enhancer binding function of FOXA1^27,28^, we found that almost half of its binding sites are characterized by high levels of the active enhancer mark H3K4me1, while the majority of binding sites of all other transcription factors did not overlap with this histone modification (**Fig. 2a, Extended Data Fig. 3a**). Interestingly, high affinity FOXA1 sites were remarkably less accessible compared to low affinity sites (**Extended Data Fig. 3a**), which is in agreement with the ability of this transcription factor to bind and subsequently open compacted chromatin, rendering it accessible to other transcription factors^25,26^. The fact that binding affinities of FOXA1 showed a strong correlation with H3K4me1 (**Extended Data Fig. 3 a-c**), indicates that this epigenetic mark is permissive for FOXA1 binding. In contrast, levels of DNA accessibility and H3K4me3 were relatively stable (and higher than for FOXA1) at binding sites of the other transcription factors over the complete range of K_d_^Apps^, indicating that these require hyper-accessible sites for binding to occur, regardless of transcription factor concentration.

Finally, we observed that FOXA1 binding at promoters is characterized by significantly lower affinities (higher numerical K_d_^Apps^) compared to binding at other regulatory elements (**Fig. 2c**). This indicates that these sites are only bound at very high FOXA1 concentrations. To validate this finding, we intersected our genome-wide FOXA1 binding affinities with previously identified^28^ sites that gain FOXA1 binding upon FOXA1 overexpression, and found that FOXA1-gained sites displayed significantly lower affinity compared to retained sites (**Fig. 2d**). In addition, sites that gain FOXA1 binding primarily consisted of promoters (75 %), while only 25 % of sites that retained pre-existing FOXA1 binding mapped to promoters (**Extended Data Fig. 4b**) and displayed higher DNA accessibility and H3K4me3 levels, but lower H3K4me1 levels (**Extended Data Fig. 4c-e**). These results demonstrate that chromatin context greatly influences binding affinities of transcription factors to DNA, thereby emphasizing the added value of investigating transcription factor binding affinities in a native chromatin context compared to naked DNA. Furthermore, for the transcription factor FOXA1, we observed an anti-correlation between binding affinities and DNA accessibility, which is the opposite from what we observed for SP1, YY1 and MYC/MAX, thus providing evidence for fundamentally different molecular interactions with the genome by different transcription factors.

### Concentration dependent target binding

Inspired by the observed dynamic binding of genomic targets at different transcription factor concentrations, we investigated the correlation between apparent binding affinities and regulation of distinct biological processes. To this end, we performed permutation-based gene set enrichment analyses to identify processes and pathways associated with specific transcription factor concentrations. For each transcription factor, we detected between 780 and 1603 significantly enriched gene sets from various databases at an FDR of 0.05 (**Fig. 3a**), revealing widespread gene regulation dynamics through transcription factor concentrations. Interestingly, gene modules that associate with high affinity transcription factor binding are often related to essential cellular processes such as energy production, translation and metabolism (**Fig. 3b-d**), which could suggest that these are ubiquitously active gene modules and only require minimal regulation. In contrast, biological processes that are specific for specialized cell types are more frequently associated with lower affinity binding sites, indicating that they can be activated in the cells that require them by increasing the transcription factor concentration towards the higher nanomolar range. In addition to biological processes, we identify hundreds of gene sets for transcription factor targets (TFT database) (**Fig. 3a**), which implies that depending on their concentration, transcription factors can cooperate with other transcription factors to activate target genes to establish complex gene regulatory networks. The most prominent example of this in our data is FOXA1, which associates with 203 transcription factor target modules. This is in line with the pioneering function of FOXA1, known to cooperate with other transcription factors to enable and regulate binding at their respective target genes^25^. Moreover, analysis of target gene sets of FOXA1 highlighted histone H3 acetylation and ATP-dependent chromatin remodelling, suggesting that positive feedback loops at the level of chromatin remodelling may stimulated by FOXA1 transcription factor activity in MCF-7. Together, these results underscore the potential of concentration dependent binding of transcription factors to their targets and subsequent activation of distinct biological processes in a concentration dependent manner.

**Figure 3.**
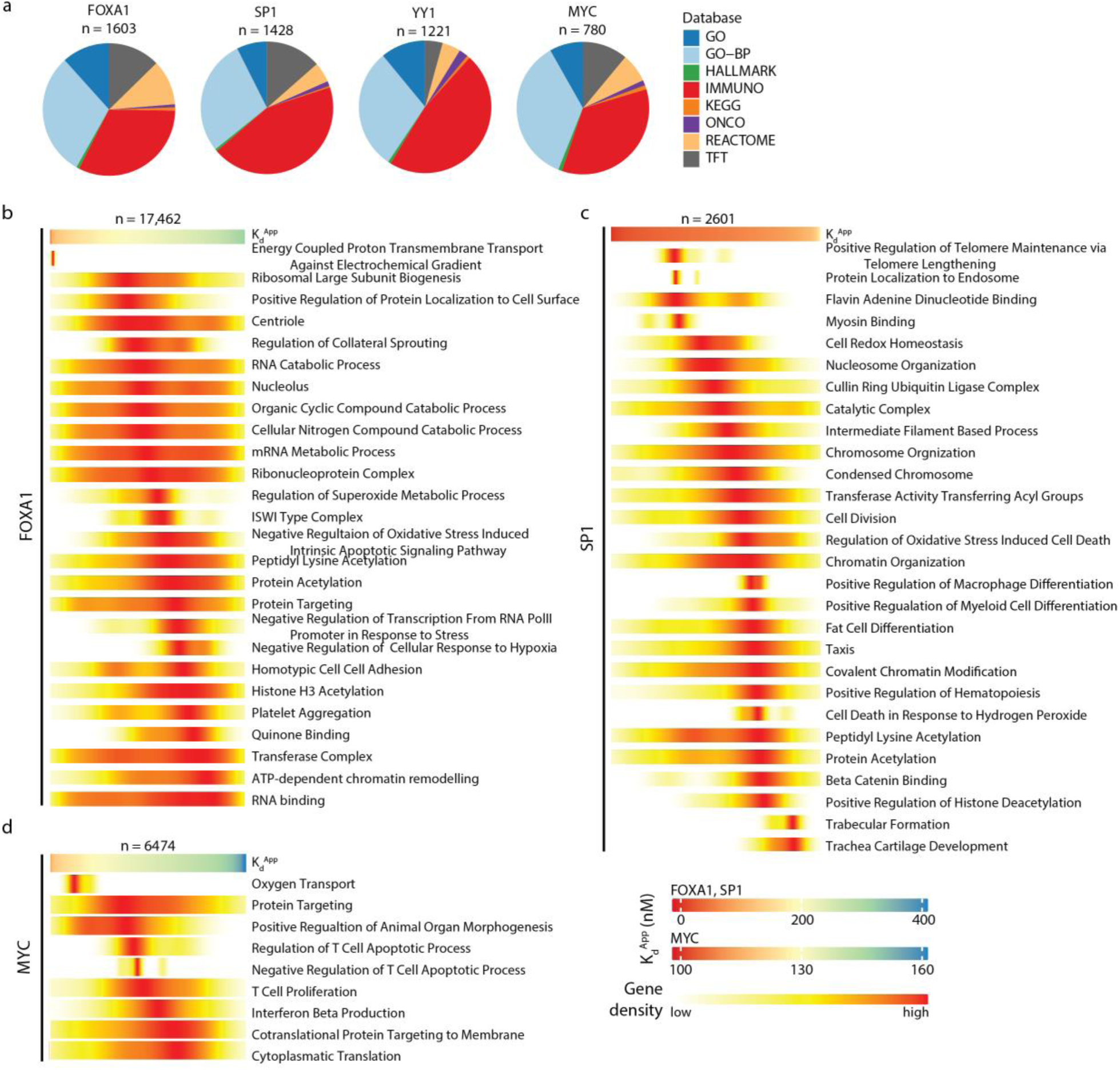
Concentration dependent binding of transcription factor target genes. (**a**) Pie charts representing the proportions of significantly enriched gene sets (FDR<0.05) per Molecular Signatures Database collection for the different transcription factors. (**b-d**) Heatmaps representing enrichment of genes from various gene sets over the range of K_d_^Apps^ for SP1, FOXA1 and MYC/MAX complex in MCF-7. Sites are ranked by K_d_^Apps^ (top heatmap per experiment) and gaussian kernel density estimates of the density of highly significant gene sets (FDR<0.001) over the ranked K_d_^Apps^ values are visualized to show that some gene sets are enriched at certain transcription factor concentrations.

### DNA sequence fine-tunes binding affinities

Given the fact that most if not all transcription factors interact with specific DNA motifs, we investigated the contribution of DNA sequences to transcription factor binding affinities by performing motif searches on high and low affinity binding sites for all transcription factors used in this study (**Extended Data Fig. 5**). Interestingly, while high affinity binding motifs were specific for the different examined transcription factors and consisted of the consensus motifs of the respective transcription factors, motifs associated with low affinity binding sites were similar in the different experiments (e.g. the common promoter CCAAT motif, which was enriched in low affinity binding sites of SP1, YY1 and MYC). This was in line with our earlier observation that low affinity binding of YY1, SP1 and MYC frequently map to the same promoter region (**Extended Data Fig. 2d**). Together, this indicates that at high expression levels, transcription factors will bind accessible chromatin regions independently of the presence of transcription factor binding motifs, underlining the dominant effect of the epigenome landscape of transcription factor binding.

Next, we hypothesized that small variations in the DNA sequence in or around consensus motifs may fine-tune apparent binding affinities. To investigate this, we made use of Castaneus/129/Sv hybrid mouse embryonic stem cells^29^ (further referred to as F121) to identify apparent binding affinity quantitative trait loci (QTLs). After aligning sequencing reads to either Castaneus or 129/Sv based on SNPs, we were able to determine allele-specific K_d_^Apps^ for 6066 sites (**Fig. 4a**). We identified 1272 QTLs that contain SNPs in the core consensus motif of YY1 (ATGG/CCAT). In the example shown in **Extended Data Fig. 6a**, we identified an almost two-fold higher apparent binding affinity in the Castaneus allele for Qars promoter, which contained two additional YY1 binding motifs in the tested site. We could validate these results by DNA affinity purifications followed by quantitative mass spectrometry, in which Yy1 showed a two-fold enrichment when bound to this sequence compared to the sequence from the 129/Sv allele (**Extended Data Fig. 6b**). However, almost 5000 sites that displayed allele-specific K ^Apps^ carry SNPs outside the core YY1 motif, indicating that for most sites, sequence variations in the YY1 motif alone does not account for the observed allele-specific variation in K_d_^Apps^. Indeed, differences in allele-specific motif-scores of the YY1 motif did not correlate with differences in allele-specific K_d_^Apps^ (**Fig. 4b**, Spearman correlation r = 0.00, p-value = 0.87; X^2^-test p-value = 0.94). However, when we further investigated sites in which SNPs were adjacent to, but not overlapping with the YY1 motif, we observed that differences in allele-specific K_d_^Apps^ were larger for sites where the SNP and YY1 motif were located close to each other (**Fig. 4c**). This finding suggests that binding of other transcription factors in the vicinity of putative YY1 binding sites influence the binding of YY1 itself. Together, this indicates that small sequence variations influence the binding of YY1 and its co-factors or interaction partners, thereby fine-tuning the binding affinity of YY1. However, this DNA sequence-based fine-tuning process is in turn subjugated to epigenetic marking at *cis-*acting regulatory regions for YY1.

**Figure 4.**
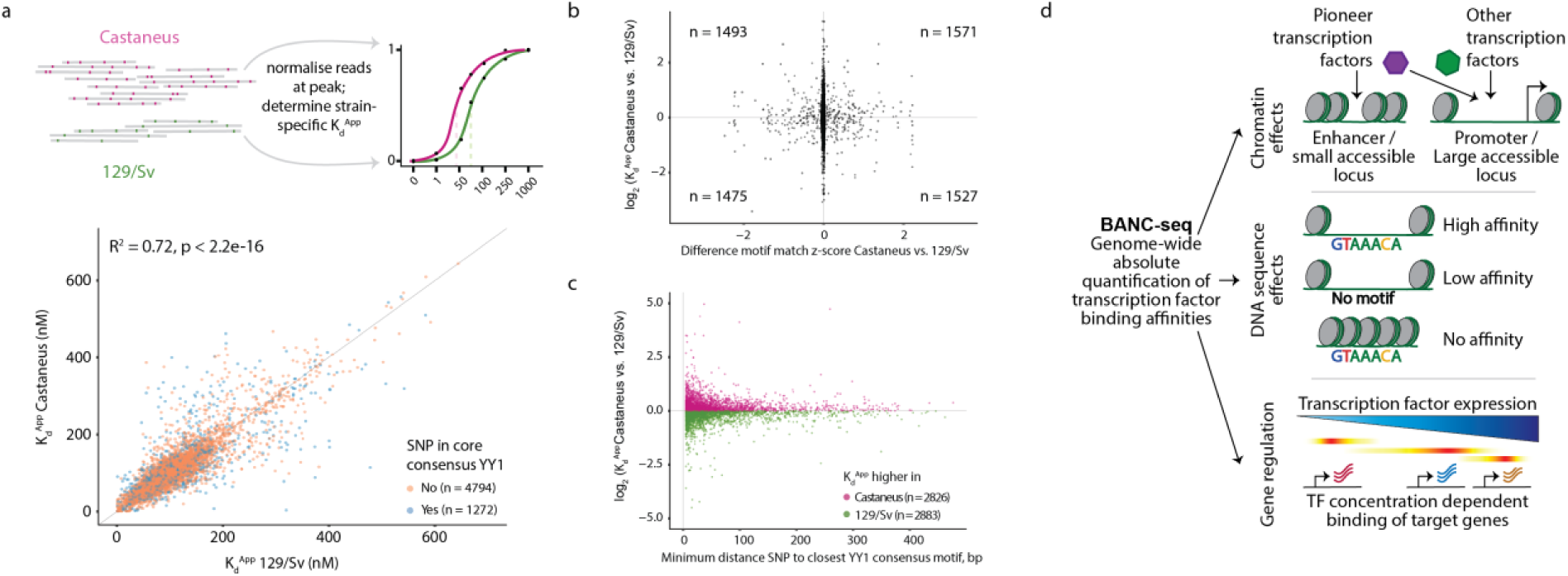
Consensus motifs are not the major determinants of transcription factor binding affinities. (**a**) Top; analysis strategy for the determination of allele-specific K_d_^Apps^. Sequencing reads were mapped to two alleles based on the presence of SNPs and separately processed further as described for the other experiments to determine allele-specific K_d_^Apps^. Bottom; scatterplot showing allele-specific K_d_^Apps^, for sites at which high confidence K_d_^Apps^ could be determined for both alleles, colour coded for whether or not a SNP was overlapping with the YY1 core consensus motif ATGG/CCAT. (**b**) Scatterplot representing the difference in allele-specific K_d_^Apps^ (y-axis) relative to the difference in alleles-specific YY1 motif-match score (x-axis) for sites at which high confidence K_d_^Apps^ could be determined for both alleles. (**c**) Scatter plot showing the difference in allele-specific K_d_^Apps^ (y-axis) relative to the smallest distance of a SNP to a YY1-core consensus motif within each site. (**d**) Summary illustrating transcription factor binding affinity regulation by the chromatin context and DNA sequence, as well as concentration dependent binding of transcription factor targets.

In summary, we have established a sequencing based method called BANC-seq that adds a quantitative dimension to genome-wide transcription factor biology. By employing BANC-seq to transcription factors FOXA1, SP1, YY1 and MYC/MAX in different cellular contexts, we show that the chromatin landscape is the major determinant for high affinity transcription factor binding, while sequence motifs appear to be secondary in the regulation of binding to DNA (**Fig. 4d**). Furthermore, we have identified K_d_^App^-range specific gene sets for different transcription factors, strongly supporting the model that the expression level of a transcription factor influences activation of larger subsets of target genes as a function of its apparent affinity range for the pre-existing chromatin environment.

## Discussion

The *in vivo* protein-DNA interaction landscape is affected by different factors including DNA sequence, chromatin state and organization, and transcription factor abundances. The BANC-seq method that we have developed here allows a quantitative assessment of the interplay between these three factors in a genome-wide manner. The ability to profile genome-wide absolute binding affinities in a native chromatin context provides a currently missing quantitative link between transcription factor expression and target gene regulation.

A key principle of the BANC-seq methodology is exogenous addition of a tagged transcription factor to permeabilized native nuclei, which also contain the endogenous transcription factor of interest. Here, the exogenous and endogenous transcription factor will compete for protein and DNA binding and in time a binding equilibrium is established in which the fraction of bound FLAG-tagged transcription factor at any given time will be steady, depending on the added concentration. Importantly, residence time for most transcription factors at their binding sites is in the order of seconds^30^, which indicates there is sufficient time to establish a binding equilibrium using our protocol. In practice this means that, as long as a binding equilibrium is established, the expression level of the endogenous transcription factor will not affect the K_d_^App^ determination.

While we show that the chromatin context has a strong impact on genome-wide transcription factor binding affinities, we cannot exclude that BANC-seq potentially underestimates binding affinities due to technical reasons. For example, the experimental conditions in which binding assays are performed greatly influence the detected apparent K_d_^Apps^. We performed our BANC-seq experiments in near physiological conditions, that is at 37°C, with high salt, protein and DNA concentrations. Indeed, performing assays at lower temperatures or salt concentrations, greatly influences observed binding affinities^20,21,31,32^. This is further illustrated by the binding affinity of MYC/MAX to the E-box motif, which reduces 20-fold when tested in phosphate-buffered saline (PBS, used in this study) compared to tris-buffered saline (TBS)^33^. Therefore, since the determined affinities are apparent to this method and possibly deviate from *in vitro* assays because we measure in complex protein mixtures, with additional unknown effectors, we refer to apparent dissociation constants (K_d_^Apps^) throughout this manuscript.

Nevertheless, it is important to evaluate previously reported low nanomolar K_d_ values in the context of the size of the genome. For example, there are over 250,000 CACGTG and 25,000,000 CCAT (MYC/MAX and YY1 binding motif, respectively) motifs in the human genome, while (depending on the nuclear volume^34^ used for the calculation) only ∼400 transcription factor molecules are expected to be present in the nucleus at a concentration of 3nM (lowest reported K_d_ for YY1^24^). If high affinity binding to consensus motifs could already occur at this concentration, MYC and YY1 would be outnumbered by the number of potential binding sites by several orders of magnitude, creating a regulatory system that would be completely dominated by stochasticity. These observations also highlight the essentiality of measuring binding affinities in the context of chromatinized DNA at near physiological conditions to obtain a fundamental understanding on how epigenome organization regulates transcription factor binding.

In summary, BANC-seq represents a powerful technology to generate quantitative maps of transcription factor binding affinities across the genome and its associated epigenetic landscape. As exemplified in this study, these maps reveal fundamental insights concerning transcription factor biology and epigenetics. In addition, BANC-seq will aid the interpretation of future studies that observe epigenome remodelling or transcription factor expression dynamics. Such data will bridge absolute transcription factor expression dynamics, as determined absolute mass spectrometry-based proteomics^35–38^, to regulatory networks and substantially improve the accuracy of predicting genome-wide transcription factor binding. The observation that gene expression patterns are regulated by distinct binding affinities at regulatory elements indicates that it is essential to incorporate BANC-seq data and epigenome profiling data in order to build relevant gene regulatory networks that accurately model gene regulation by a transcription factor.

## Methods

### Cell culture

Human MCF-7 cells were grown in DMEM (Gibco) supplemented with 10% fetal bovine serum (GE Healthcare Life Sciences) and 1X penicillin–streptomycin (Gibco). F121 mouse embryonic stem cells were grown on 0.15% gelatin coated dishes in DMEM (Gibco) supplemented with 15 % fetal bovine serum (GE Healthcare Life Sciences), 5 μM beta mercaptoethanol (Sigma), 1X non-essential amino acids (Lonza), 1X Glutamax, 1 mM sodium pyruvate (Gibco), 10 mM Hepes, 1X penicillin–streptomycin (Gibco) and in-house generated Leukemia Inhibitory Factor (LIF). At ∼80% confluency, the cells were washed once with PBS, scraped from the plate, collected and washed twice with cold PBS, after which they were aliquoted and cryopreserved in HyClone fetal bovine serum with 10 % DMSO at -80°C until further processing.

For DNA pulldown followed by mass spectrometry, we harvested cells and prepared nuclear extracts as described previously^39^.

### Transcription factor binding

Cryopreserved cells were thawed quickly at 37°C and washed twice with ice-cold PBS. All subsequent steps were performed on ice unless stated otherwise. Cell membranes were lysed by adding 900 μl of ice-cold hypotonic lysis buffer containing 10 mM Tris/HCl at pH 7.5, 10 mM NaCl, 3 mM MgCl_2_ and 0.1 % IGEPAL ca-630. Nuclei were isolated by pipetting up and down 20 times, followed by centrifugation at 250 x g for 10 minutes at 4°C. Isolated nuclei were resuspended and counted in ice-cold PBS, to aliquot 2 × 10^6^ or 2.5 × 10^5^ nuclei per concentration in separate tubes for either ChIP-based or CUT&RUN-based follow up, respectively. Each tube with nuclei was resuspended in the following ice-cold incubation buffer to a volume of 20 μl: 1 % BSA, 1 mM CaCl_2_, 5 mM MgCl_2_, 1 μM ZnCl_2_, 0.1 % IGEPAL ca-630, 1× EDTA-free protease inhibitor cocktail (cOmplete, Roche), 1× PBS, supplemented with protein-of-interest at a designated concentration in diluted protein storage buffer. Recombinant FLAG-YY1 (Active Motif, Cat. nr. 81119), FLAG-MYC/MAX (Active Motif, Cat. nr. 81087), FLAG-SP1 (Active Motif, Cat. nr. 81181) or FLAG-FOXA1 (OriGene Technologies Inc, Cat. nr. TP306045), were diluted to the highest tested concentration with ddH_2_O, and further diluted in protein storage buffer of the respective supplier to ensure that the buffer conditions of all concentrations were identical. To improve complete nuclear permeabilization and diffusion of the protein of interest into the nuclei, the nuclei were briefly sonicated for 5 seconds at 4°C in a Bioruptor Pico sonicator (Diagenode). Permeabilized nuclei were incubated for 10 minutes at 37°C in a thermoshaker, rocking at 1000 rpm.

### Chromatin immunoprecipitation (ChIP)-based follow up

While shaking the nuclei at 37°C, chromatin was cross-linked by adding 1% formaldehyde to a final concentration of 1% (v/v) followed by incubation for 4 minutes and quenching by adding 0.1 volumes of 1.25 M glycine. Chromatin was recovered with an ethanol precipitation with 0.1 volumes of sodium acetate and 3 volumes of ice-cold ethanol at -20°C for 15 minutes, followed by ten minutes centrifugation at max speed. After washing once with ice-cold 70% ethanol, the purified chromatin was dissolved by shaking at 37°C in the following sonication buffer: 20mM Hepes at pH 7.6, 1% SDS and 0.25× EDTA-free protease inhibitor cocktail (cOmplete, Roche). Chromatin was sheared in a Bioruptor Pico sonicator (Diagenode) at 4°C by 5 cycles of 30 s ON, 30 s OFF.

A ChIP master mix was added to each chromatin sample to achieve the following final conditions for each ChIP reaction: 0.1% BSA, 1 x EDTA-free protease inhibitor cocktail (cOmplete, Roche), 1 μg anti-Flag antibody (clone M2, Sigma), 1 μg anti-H2Av antibody (spike-in antibody, Active Motif), 10 ng spike-in chromatin (*D*.*melanogaster*, Active motif) and 1× incubation buffer (0.15% SDS, 1% TritonX-100, 150mM NaCl, 1mM EDTA, 0.5mM EGTA, 20mM HEPES). ChIP reactions were incubated by rotating overnight at 4 °C. To each sample, 15 μl of a 1:1 mix of Protein A and Protein G Dynabead (Invitrogen) was added followed by a 90 minute incubation at 4 °C. On ice, the beads were washed 2× with Wash Buffer 1 (0.1% SDS, 0.1% sodium deoxycholate, 1% Triton, 150 mM NaCl, 1 mM EDTA, 0.5 mM EGTA, and 20 mM HEPES), 1× with wash buffer 2 (wash buffer 1 with 500 mM NaCl), 1× with wash buffer 3 (250 mM LiCl, 0.5% sodium deoxycholate, 0.5% NP-40, 1 mM EDTA, 0.5 mM EGTA, and 20 mM HEPES), and 2× with wash buffer 4 (1 mM EDTA, 0.5 mM EGTA, and 20 mM HEPES). After washing, chromatin was eluted from the beads at room temperature by incubating them for 20 minutes in a thermoshaker at 1400 rpm in 100 μl of the following buffer: 1% SDS, 0.1 M NaHCO3. The supernatant was decrosslinked overnight at 65°C in a thermoshaker at 1000 rpm by adding 20 μg of proteinase K and 4 μl of 5M NaCl. Decrosslinked DNA was purified by column purification (Zymo) and used for quantitative reverse transcription PCR (qPCR; hPTBP1 promoter: GTTTCCTGCCCGACTCCAAGAT and GAGGGGGAGAAAATGGGATCACG; gene desert: AACTGGCTAGTAAGGAGTGAATG and GGGAATGGAAAGAAGTCCACTAT) or next-generation sequencing sample preparation.

### CUT&RUN-based follow up

Nuclei were placed on ice immediately after the 10-minute incubation with the transcription factor of interest. Ice-cold wash buffer 1 (150 mM NaCl, 20 mM Hepes pH 7.5, 0.5 mM Spermidine, 0.2 mM TritonX-100, 2 mM EDTA pH 8, 1 x EDTA-free protease inhibitor cocktail) containing 1 μg anti-Flag antibody (clone M2, Sigma) was added to the nuclei to a total volume of 400 μl and incubated in a rotation wheel overnight at 4 °C. Then, inspired by the original CUT&RUN protocol^5^, each sample was washed 2x with 300 μl of ice-cold wash buffer 2 (150 mM NaCl, 20 mM Hepes pH 7.5, 0.5 mM Spermidine, 0.2 mM TritonX-100, 1 x EDTA-free protease inhibitor cocktail), followed by incubation with 150 μl of wash buffer 2, supplemented with 1.5 μl in-house generated recombinant pAG-MNase (diluted in wash buffer 2 to the working concentration), rotating at 4 °C for one hour. Samples were washed again 2x with wash buffer 2, and MNase was activated by adding 100 μl of wash buffer 3 (150 mM NaCl, 20 mM Hepes pH 7.5, 0.5 mM Spermidine, 0.2 mM TritomX-100, 2 mM CaCl_2_). The digestion reaction was performed for 30 minutes at 4 °C, and stopped by adding 100 μl 2X Stop buffer (final concentration 170 mM NaCl, 10 mM EDTA, 2.5 mM EGTA, 0.025 % digitonin), supplemented with 15 pg spike-in DNA per sample (*S*.*cerevisae*, Cell Signaling Technologies), with exception of the YY1 experiment in MCF-7. Samples were incubated at 37 °C for 30 minutes to release digested DNA fragments, followed by max speed centrifugation at 4 °C for 5 minutes and DNA purification by column purification (Zymo) prior to sample preparation for next-generation sequencing.

### Library preparation and sequencing

BANC-seq libraries were prepared using the Kapa Hyper Prep Kit (Kapa Biosystems) according to manufacturer’s protocol, with the following modifications. All input material after column purification was used to prepare libraries and depending in the concentration of the starting material, 2.5 μl or 5 μl of the NEXTflex adapter stock (600nM, Bioo Scientific) was used for adapter ligation of each sample. Libraries were amplified with 12 PCR cycles, followed up with one reverse and a double post-amplification clean-up was used to ensure proper removal of adapters. Samples were analyzed for purity using a High Sensitivity DNA Chip on a Bioanalyzer 2100 system (Agilent). Libraries were paired-end sequenced on an Illumina NextSeq500.

### Mass spectrometry based whole cell or nucleus proteomics

Sample pellets from 2 million cells were taken before and after the hypotonic lysis step in the nuclear isolation procedure. Pellets were dissolved in 50 uL lysis buffer (4 % SDS, 0.1 M Hepes at pH 7.6 and 0.1 M DTT) by boiling at 95°C for 5 minutes followed by 5 sonication cycles of 30 s ON / 30 s OFF on the Bioruptor Pico sonicator (Diagenode) at 4°C. We added 1.25ug Universal Proteomics Standard-2 (Sigma) spike-in to each sample for absolute copy number quantification per cell and nucleus. Proteins were digested and cleaned for mass spectrometry analysis using the filter assisted sample preparation (FASP^40^). In short, dissolved proteins were denatured in 8 M urea, loaded on a 30 kDa filter, and alkylated with 50 mM iodoacetic acid. The filter was washed three times with urea buffer and three times with 50 mM ammonium bicarbonate buffer, followed by overnight trypsin digestion (Promega) and collection in ammonium bicarbonate buffer. Peptides were acidified with trifluoroacetic acid and desalted on C18 (Empore) StageTips^41^. For each sample, peptides were separated on an online Easy-nLC 1000 (Thermo Scientific) using a 4-minute (7 % to 9 %) acetonitrile gradient, followed by a 214 min gradient of acetonitrile (9 % to 32 %), followed by washes at 50 % and 95 % acetonitrile for 240 min of total data collection. Mass spectra between 350 to 1300 m/z were collected on a Q-Exactive HFX mass spectrometer (Thermo Scientific) in Top20 mode with a full-MS and dd-MS^2^ resolution of 120,000 and 15,000, respectively. Acquired mass spectra were analyzed with MaxQuant 1.6.0.1^42^ with default settings and by searching the human Uniprot protein database downloaded in June 1017, and with the spike-in protein database supplied by the manufacturer. Intensity based absolute quantification was performed as described in^35^, by extrapolating the absolute abundance of each protein from a linear regression between the log-transformed iBAQ^43^ values and the log-transformed concentrations of the spike-in proteins. Detected transcription factor proteins were identified with the TFCheckpoint database^44^, version ‘TFCheckpoint_download_180515’, by selecting proteins in the class ‘TFclass’.

### Sequence data processing, quantification and spike-in normalisation

Pre-processing of generated sequencing data was performed automatically with workflow tool seq2science v0.5.4^45^. Briefly, paired-end reads were trimmed with fastp v0.20.1 with default options. Reads were aligned to the hg38 or mm10 reference genome, as well as to the respective spike-in genome for each experiment (S.cerevisiae-74-D694-2.0, dm6 or E.coli_C142) with bwa-mem2 v2.1 with options ‘-M’. Mapped reads were removed if they did not have a minimum mapping quality of 30, were a (secondary) multimapper or aligned inside the ENCODE blacklist. Afterwards, duplicate reads were removed with Picard MarkDuplicates v2.23.8. Peaks were called with macs2 v2.2.7 with options ‘--keep-dup 1 --buffer-size 10000 --call-summits’ in BAMPE mode. Samtools v1.9 was used to quantify total read counts per sample for target (hg38 or mm10) and designated spike-in (S.cerevisiae-74-D694-2.0, dm6 or E.coli_C142) genome. Subsequent analyses were performed in R (v 4.0.3) and data was visualised with the ggplot2 and ComplexHeatmap^46^ packages, unless stated otherwise.

For read quantification at transcription factor binding sites, we used the peak locations of the sample with the highest transcription factor concentration. To prevent peak size bias in downstream analyses we trimmed peaks to the median peak length of all peaks around the peak summit. For samples of each concentration we used featureCounts^47^ v1.6.3 to count reads per peak (with options -p -C -O -g GeneID -s 0 -F SAF -B), and normalized raw counts to reads per million spike-in reads. In addition, we used bedtools to generate bigwig files that represent the spike-in normalized signal of BANC-seq samples to be visualized in the UCSC genome browser.

### K_d_^Apps^ determination

Transcription factor binding parameters were calculated with a similar approach as we reported previously for proteomics workflows^7^. For every peak, we scaled the normalized read counts to the sample with the highest signal. These relative fractions bound over transcription factor concentration was fitted to the following Hill-like curve:

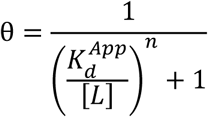

In this formula θ represents the observed relative fraction of bound transcription factor, *[L]* represents the known transcription factor concentration, *K*_*d*_^*App*^ is the apparent dissociation constant describing the concentration at which half of the DNA is bound by the transcription factor, and *n* is he Hill coefficient describing the rate at which binding saturates. The unknown parameters *n* and *K*_*d*_^*App*^ were fit using non-linear least squares regression with the nlsLM function in the minpack.lm R package^48^. Starting values of *n* and *K*_*d*_^*App*^ were set to 1 and a maximum of 50 iterations were performed. For follow-up analyses and visualisations, we included high confidence sites only (p < 0.01, r > 0.9).

### Allele-specific K_d_^App^ analysis

To determine allele-specific K_d_^Apps^ we used information on SNP locations from either 129S1/SvImJ (hereafter referred to as 129/Sv) or CAST/EiJ (hereafter referred to as Castaneus) and the GATK^49^ tool *FastaAlternateReferenceMaker* to create two references genomes based on mm10, in which each base overlapping with a SNP was replaced by the alternative base. Sequencing data from the experiment performed with F121 mESC were then processed for both references with the seq2science tool as described above. Before quantifying reads at peak locations, we removed reads (and their respective read mate) that had at least one mismatch to retain only reads perfectly aligning to either new reference with samtools.

### Peak annotation

The predicted regulatory function of identified binding sites was inferred by overlapping them with regulatory features in the Ensembl Regulatory Build^50^ of version GRCh38.MCF_7.20210107 for MCF-7 experiments and GRCm39.ES_Bruce4_embryonic.20201021 for F121 experiments. To overlap the latter, we first converted the mm10 mapped peak coordinates to mm39 via the UCSC liftOver tool. If a peak overlapped with more than one regulatory feature, it was assigned the one feature that had the highest rank in the order of ‘Promoter, Enhancer, Promoter flanking region, Open chromatin region, CTCF binding site, Transcription factor binding site’. To detect overlap between high and low affinity assigned promoters per transcription factor, we first combined YY1 bound sites from both the ChIP-seq and CUT&RUN-based follow up experiment. Then, we selected promoters only, and separately for the 20% lowest and highest affinity promoters we used the intervene^51^ tool to generate upset plots.

### Motif analysis

To identify enriched motifs in high- and low affinity binding sites, we used the binding sites with the 500 lowest and highest K_d_^Apps^ for each experiment as input for gimme motifs^52^ and selected the top enriched motifs per transcription factor in either high or low affinity binding sites for visualisation. To assess the effect of motif perturbations on K_d_^Apps^, we determined the YY1 motif match score in both alleles of the hybrid mESCs with the scan function of gimme motifs and combined the differences in allele-specific K_d_^Apps^ with differences in allele-specific motif match scores. Finally, we determined the minimal distances between consensus motifs and SNPs in each binding site to combine this with differences in allele-specific K_d_^Apps^, and used the SNP location closest to YY1 motifs (+/-50 bp) as input for gimme motifs to find motifs in the vicinity of these SNPs.

### Concentration dependent target binding

We used GREAT^53^ to assign each identified binding site to a single gene whose transcription start site is the closest within 1,000,000 bp. We used the GO, GO-BP, IMMUNO, TFT, ONCO, KEGG and REACTOME data sets from MSigDB v7.2^54^ for gene set enrichment analysis. To identify gene sets that associate with a certain transcription factor concentration, we tested if K_d_^Apps^ assigned to the genes within each gene set vary significantly less compared to random. To this end, we calculated the coefficient of variation of K_d_^Apps^ that could be assigned to genes within each gene set. Next, we shuffled the K_d_^Apps^ values 100,000 times and determined how often the actual coefficient of variation would be smaller than by chance, to determine a permutation based false discovery rate.

### Integrating epigenome data

To integrate information on genome-wide transcription factor binding affinity with epigenome data, we downloaded hg38-mapped MCF-7 as well as mm10-mapped mESC ChIP-seq and ATAC-seq data from the ENCODE-portal. When more than one experiment for the same bait was available, we selected the experiment that had most reads sequenced. Samples that were treated with drugs or genetically modified were excluded from integration. Aligned reads were quantified with featureCounts at either transcription factor binding sites or random genomic loci, normalised to reads per million mapped reads and the average signal of 5 control samples was used to compute fold changes over control. Normalised signal of ATAC-seq or ChIP-seq data at transcription factor binding sites for which we could define high confidence K_d_^Apps^ were visualized with ComplexHeatmap to visualize the histone landscape alongside different regulatory features and their associated K_d_^Apps^. In addition, we divided high confidence sites into quintiles based on K_d_^App^, and visualized the epigenome data per quintile for each experiment as boxplots. Rho (r) and p-value from Spearman correlation of the respective epigenome signal and K_d_^Apps^ are included in the boxplots.

### Overlap with endogenous ChIP-seq data

To determine the overlap between binding sites of FLAG-tagged transcription factors in BANC-seq with endogenous transcription factor binding sites, we retrieved hg38-mapped ChIP-seq data from the ENCODE-portal for SP1, FOXA1, MYC and YY1. For FOXA1, we also included peaks identified after FOXA1 overexpression^28^. For overlap of FLAG-YY1 with endogenous mYy1 binding sites in mESCs, we computed the overlap with one ChIP-seq sample^19^, which we downloaded and processed with seq2science to define Yy1 binding peaks. For all data sets and experiments, bedtools was used to identify peaks that overlapped between our experiments and endogenous ChIP-seq peaks.

### DNA pulldown followed by mass spectrometry

To validate sequence specific binding of Yy1, a DNA pulldown was performed as described previously^39^ with nuclear extracts of F121 mESCs and the following DNA oligos: Cast_Fw: TCCTATTGGTCCATGAGCAAAGGTCGCTGTTCAGATGGGGCCCAAAGT, Cast_Rv: ACTTTGGGCCCCATCTGAACAGCGACCTTTGCTCATGGACCAATAGGA, 129Sv_Fw: TCCTATTGGTCAATGAGCAAAGGTCGCTGTTCAGATGAGGCCCAAAGT, 129Sv_Rv: ACTTTGGGCCTCATCTGAACAGCGACCTTTGCTCATTGACCAATAGGA. DNA oligos were ordered via custom synthesis from Integrated DNA Technologies with 5’ biotinylation of the forward strand and annealed using a 1.5 X molar excess of the reverse strand. DNA affinity purifications were performed as described previously^39^. In short, 500 pmol of DNA oligonucleotides were immobilized using 20 μl of Streptavidin-Sepharose bead slurry (GE Healthcare, Chicago, IL). Then, 500 μg of nuclear extract and 10 μg of non-specific competitor DNA (5 μg polydAdT, 5 μg polydIdC) were added to each pulldown. After extensive washing, samples were prepared for mass spectrometry analysis or western blotting. For mass spectrometry analysis, beads were resuspended in elution buffer (2 M urea, 100 mM TRIS (pH8), 10 mM DTT) and alkylated with 50 mM iodoacetamide. Proteins were digested on beads with 0.25 μg of trypsin for 2 hours. After elution of peptides from beads, an additional 0.1 μg of trypsin was added and digestion was continued overnight. Peptides were labelled on Stage tips using dimethyl labelling as described previously^39^. Each pulldown was performed in duplicate and label swapping was performed between duplicates to avoid labelling bias. Matching light and heavy peptides were combined and analysed on an Orbitrap Exploris (Thermo) mass spectrometer with acquisition settings described previously^55^. RAW mass spectrometry data were analysed with MaxQuant 1.6.0.1 by searching against the UniProt curated mouse proteome (released June 2017) with standard settings. Protein ratios obtained from MaxQuant were used for outlier calling. An outlier cut-off of 1.5 inter-quartile ranges in two out of two replicates was used and results were visualized with python.

## Acknowledgements

We thank Matthew M. Makowski, Guido van Mierlo and all members of the Vermeulen laboratory for fruitful discussions. We thank the laboratory of Joost Gribnau for sharing the hybrid mouse embryonic stem cells for this study. We thank Samy Kefalopoulou and Peter Zeller of the Hubrecht Institute for technical support with the CUT&RUN protocol. The Vermeulen laboratory is part of the Oncode Institute which is partly financed by the Dutch Cancer Society (KWF). Furthermore, work in the Vermeulen laboratory is supported by the gravitation program CancerGenomics.nl from the Netherlands Organization for Scientific Research (NWO), and by an ERC Consolidator Grant (SysOrganoid;771059).

## Author Contributions

R.G.H.L. and M.V. conceived the study. R.G.H.L. designed the methodology and analyses. H.K.N. adapted the methodology to the CUT&RUN based follow-up. R.G.H.L. and H.K.N. performed BANC-seq experiments and analysed the data. M.P.B. and L.A.L. prepared the sequencing libraries and performed next generation sequencing. P.W.T.C.J. and C.G. performed mass spec experiments. H.K.N., R.G.H.L., C.G., S.J.v.H., C.L., S.A.T. and M.V. edited the manuscript.

## Supplementary Information

is available for this paper.

## Correspondence

to Rik Lindeboom or Michiel Vermeulen.

## Data availability

Next-generation sequencing data have been deposited to the Gene Expression Omnibus (GEO) with accession code GSEXXXXXX.

## Code availability

Data analysis script to perform K_d_^App^ determination from count files are available on https://github.com/HNeikes/BANCseq.

**Extended Data Figure 1.**
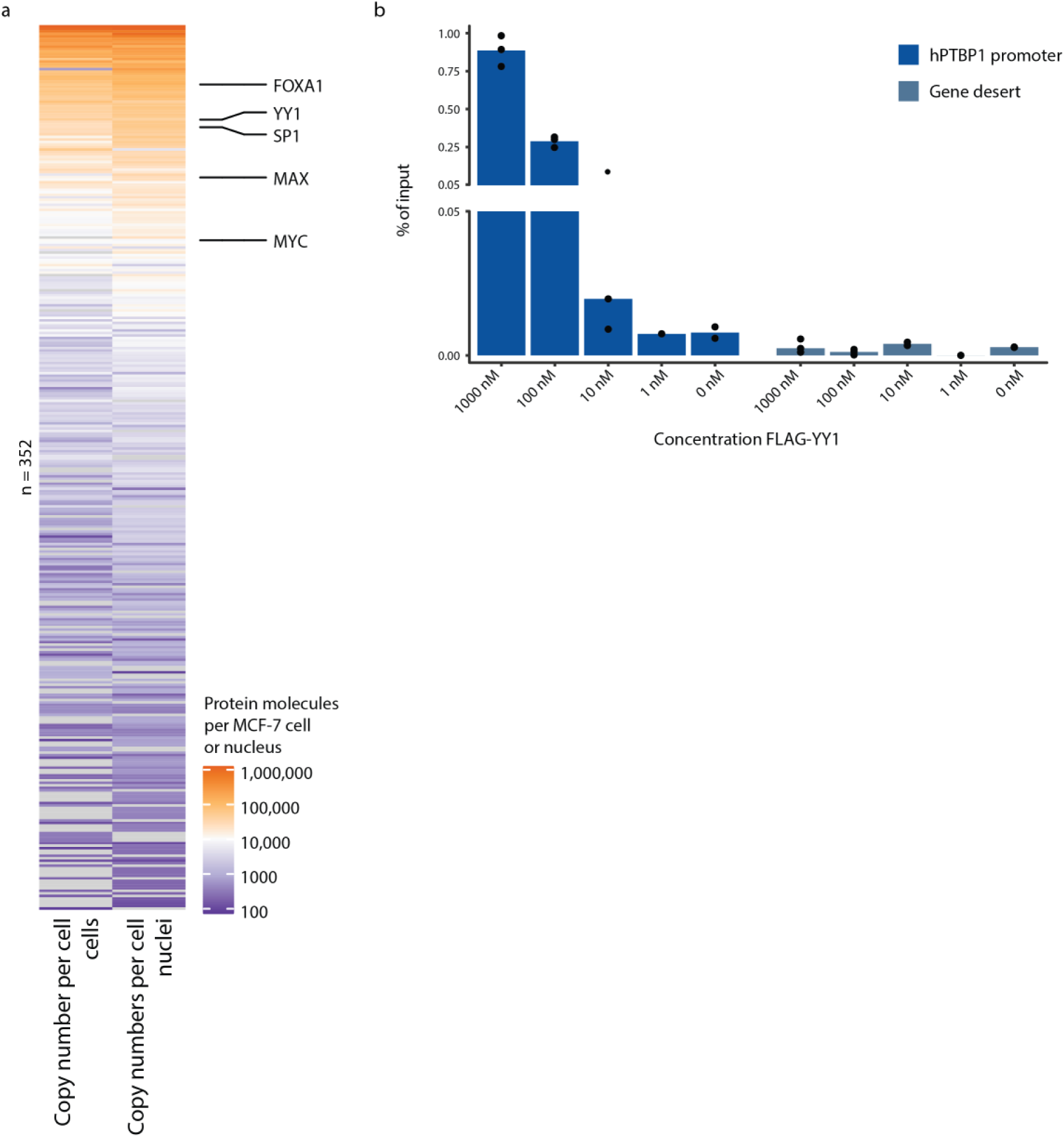
Technical support for the pilot BANC-seq experiment. (**a**) Heatmap showing copy numbers per cell or nucleus of detected transcription factors before and after nuclear isolation. (**b**) Recovery (as percentage (%) of input chromatin) at the human PTBP1 promoter and a random genomic site by ChIP-qPCR per titration point of FLAG-YY1 in MCF-7 cells.

**Extended Data Figure 2.**
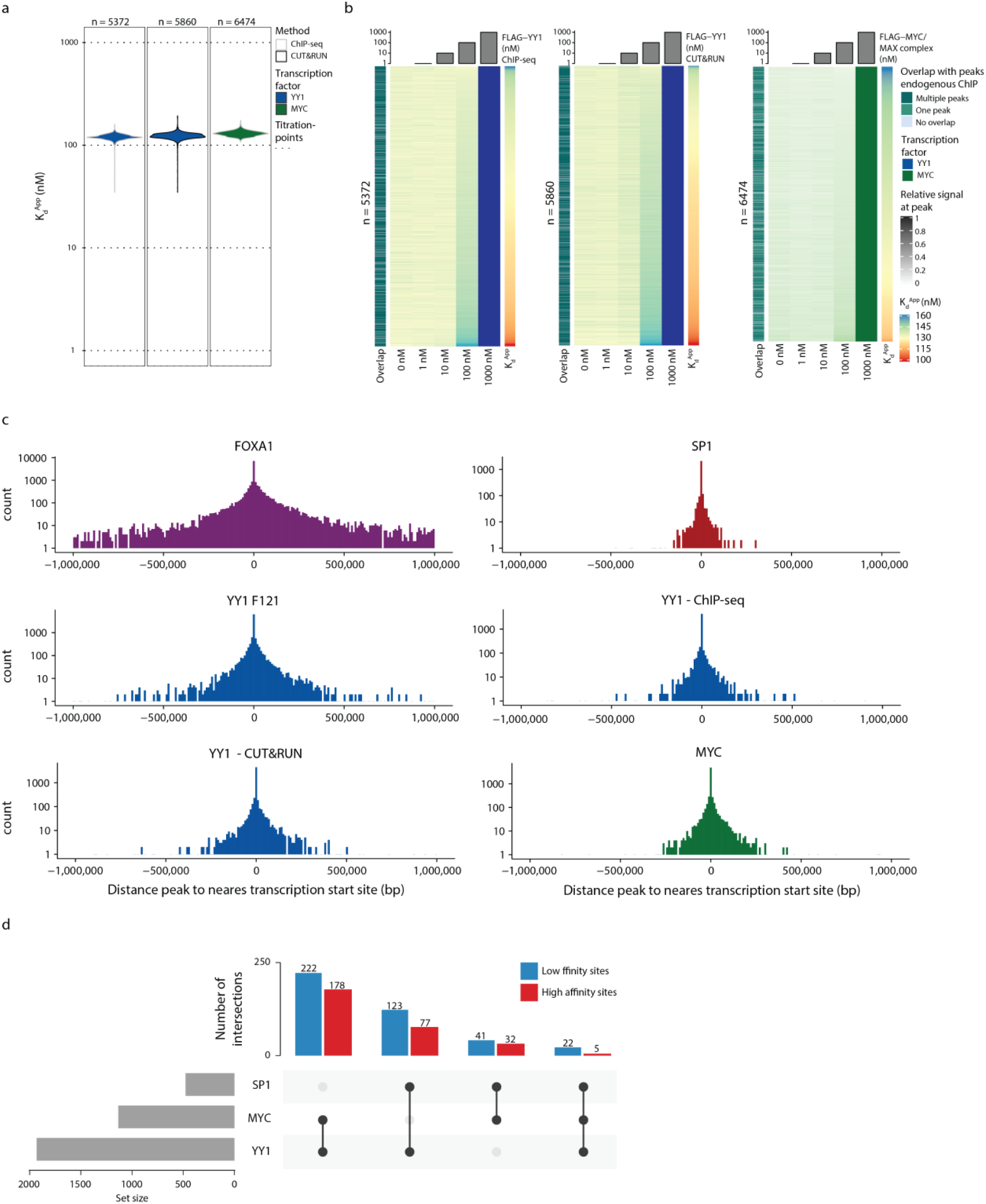
Overview of results of additional BANC-seq experiments. (**a**) Distribution of K_d_^Apps^ of c-MYC and YY1 in MCF-7 cells. YY1 apparent binding affinities were probed either by ChIP-seq or CUT&RUN-based follow up of the protocol. Dotted lines indicate the tested concentrations per experiment. (**b**) Heatmap representing spike-in normalised sequencing reads relative to the highest signal for the same experiments as in (a). Each row represents one transcription factor binding site. The overlap of each binding site with peaks from endogenous ChIP-seq experiments of the same transcription factor is shown to the left of each heatmap, while K_d_^Apps^ to the right. (**c**) Distance (bp) of identified transcription factor binding sites relative to the nearest transcription start site (TSS). (**d**) Blue and red barplot representing the overlap between promoters bound by YY1, MYC or SP1, separately for promoters assigned to be 20% highest or lowest affinity binding sites for all possible combinations of the three transcription factors. Grey barplot to the left representing the total size of each promoter set.

**Extended Data Figure 3.**
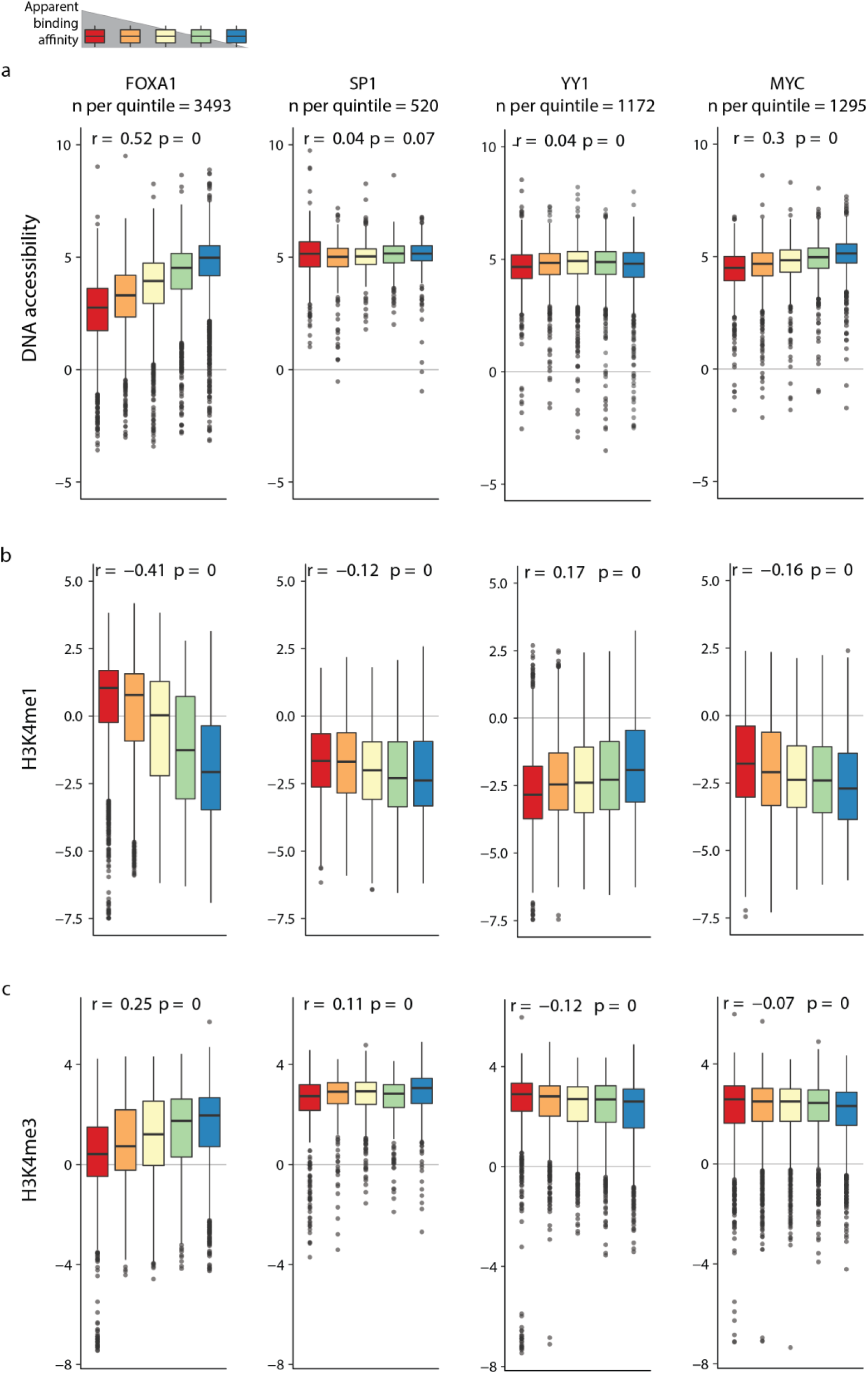
Overview of the chromatin context and correlation with K_d_^Apps^ for all transcription factors. Boxplots representing log_2_ fold change of ATAC-seq (**a**), H3K4me1 ChIP-seq (**b**) or H3K4me3 ChIP-seq (**c**) signal over the mean signal in matched control tracks for all tested transcription factors at sites with high confidence K_d_^Apps^ fitted. Values are ranked by K_d_^App^ and divided into quintiles based on K_d_^Apps^ per experiment. Rho (r) and p-value from Spearman correlation of the respective epigenome signal and KdApps are included in the boxplots. Box plots were drawn with the center line as the median, the hinges as the first and third quartiles, and with the whiskers extending to the lowest and highest values that were within 1.5 × interquartile range..

**Extended Data Figure 4.**
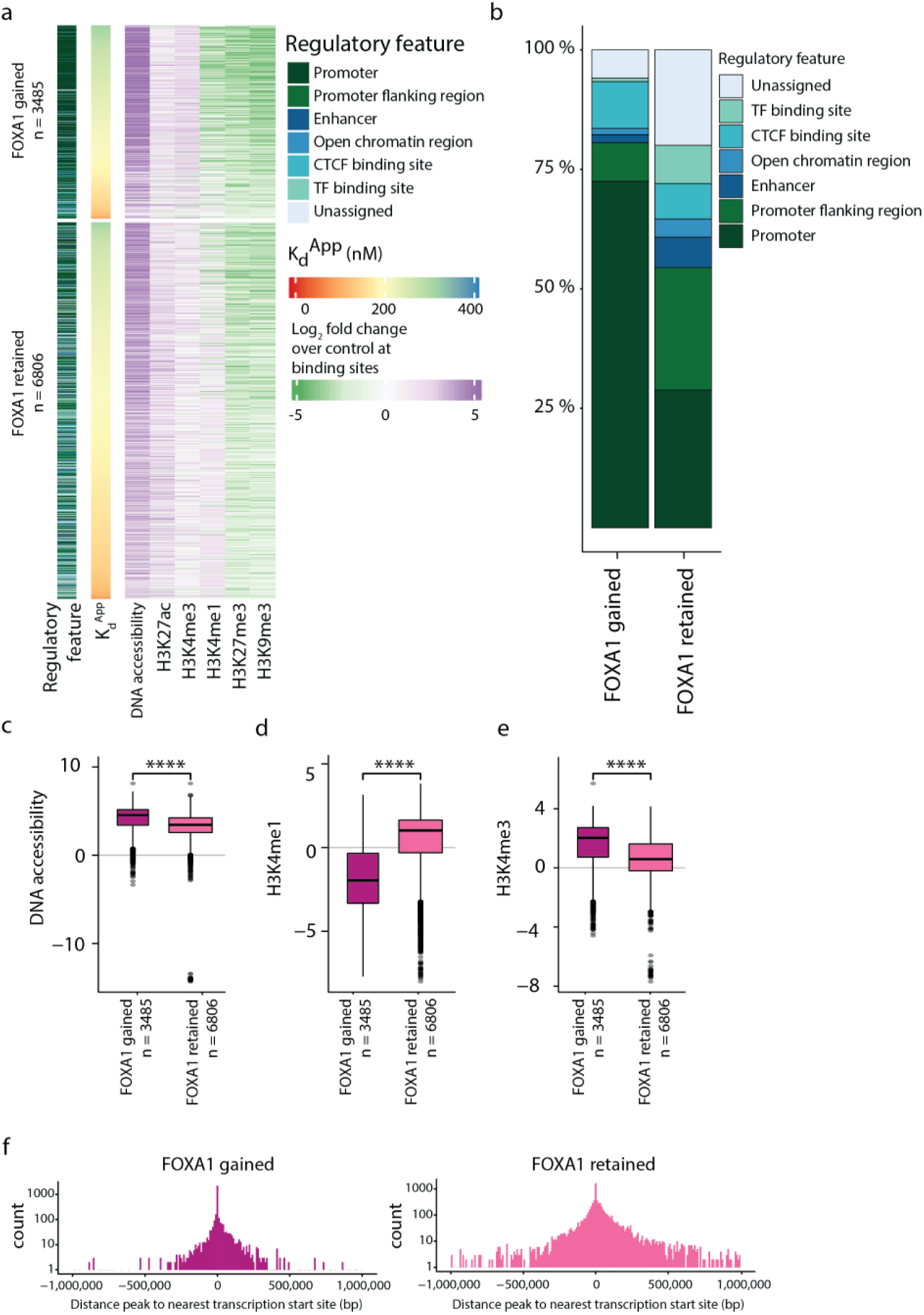
FOXA1 binds hyperaccessible promoters with low affinity upon overexpression in MCF-7. (**a**) Heatmap showing the matched epigenome dynamics at sites with high-confidence K_d_^Apps^ fitted for FOXA1 at either gained or retained sites after FOXA1 overexpression. Signal of ChIP-seq and ATAC-seq tracks for MCF-7 is shown as log_2_ fold change over the mean signal in all matched control tracks, sites are ranked by apparent binding affinity (second column), and assigned regulatory features are depicted in the first column. (**b**) Overlap of gained or retained FOXA1 binding sites with known regulatory features. (**c - e**) log_2_ fold change of ATAC-seq, H3K4me1 ChIP-seq or H3K4me3 ChIP-seq signal over the mean signal in matched control tracks, separated by sites being gained and retained sites after FOXA1 overexpression. * are used to indicate significance according to a two-sided Wilcoxon test (****: p <= 0.0001). (**f**) Distance (bp) of gained or retained sites to the nearest transcription start site (TSS). Box plots were drawn with the center line as the median, the hinges as the first and third quartiles, and with the whiskers extending to the lowest and highest values that were within 1.5 × interquartile range.

**Extended Data Figure 5.**
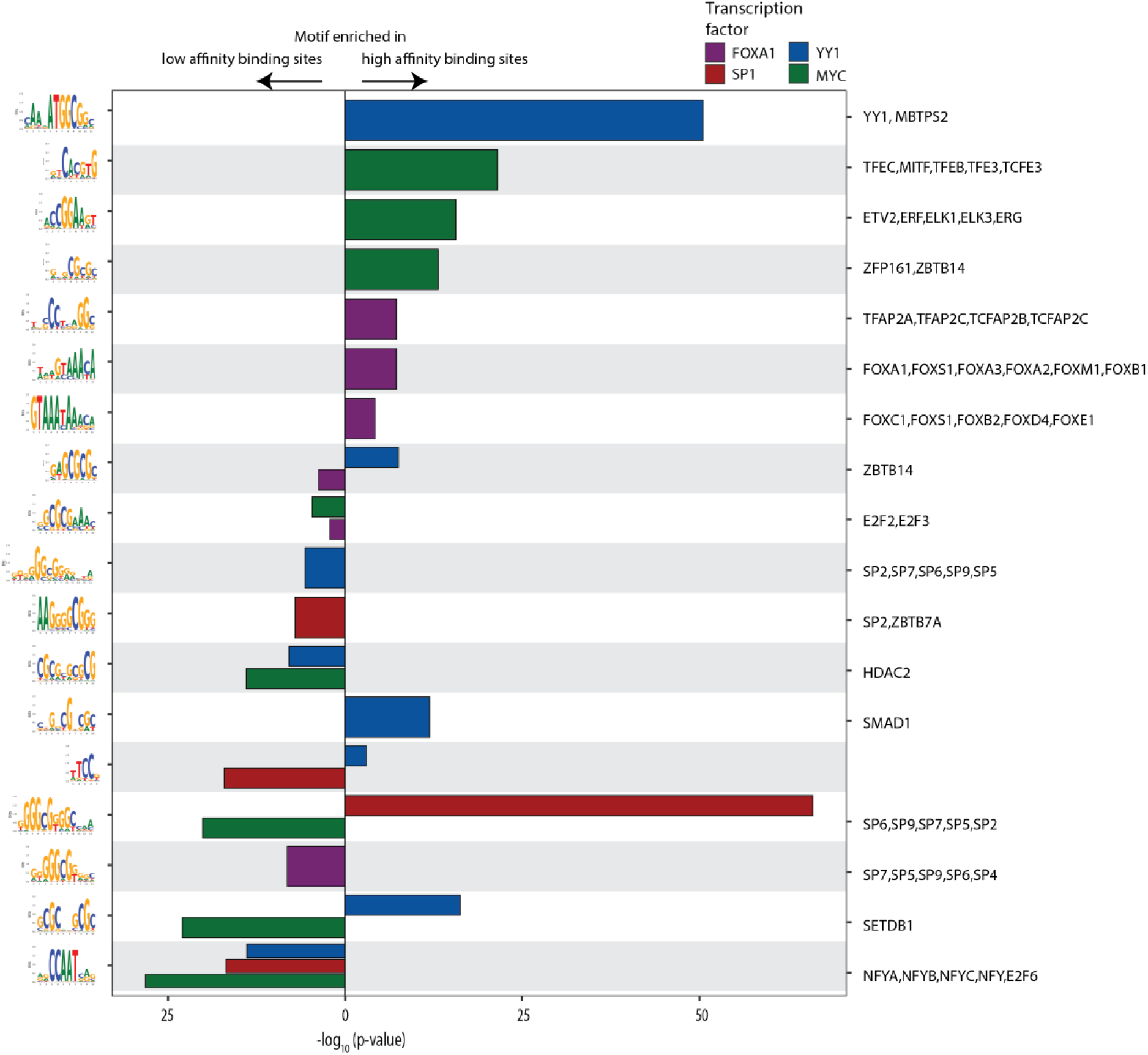
Transcription factor specific motifs versus generic motifs in high versus low affinity binding sites. Bar plot representing p-vales (-log_10_) of top motifs per transcription factor for either high or low affinity binding sites. Motif logos on the left of the plot, names of associated transcription factors (if known) to the right..

**Extended Data Figure 6.**
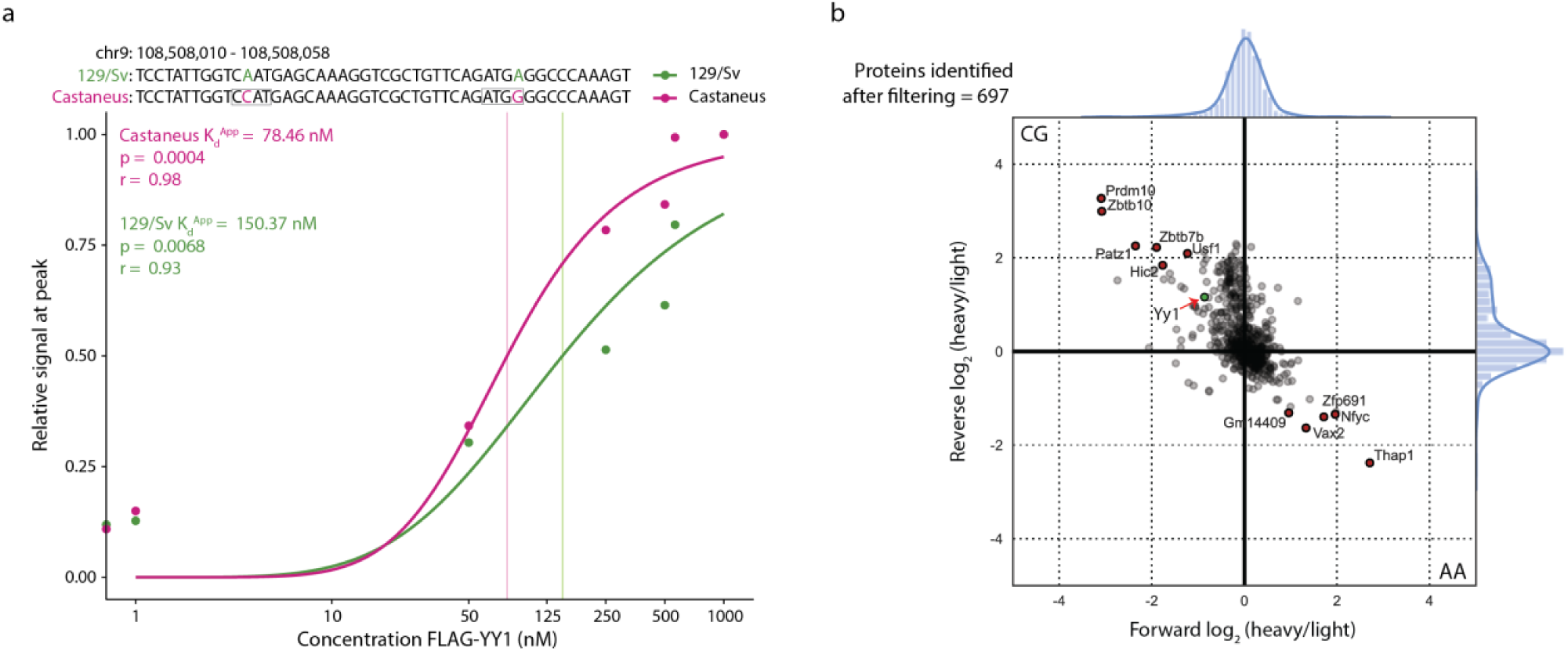
Minor sequence variations in and near the consensus motif of YY1 fine-tune apparent binding affinities. (**a**) Spike-in normalised sequencing reads per allele and titration point of FLAG-YY1 in F121 mESCs relative to the highest signal at the *Qars* promoter. (**b**) Binding ratios (log_2_ scale) of proteins identified by DNA-pulldown followed by mass spec with oligos identical to the sequences in (a) Red arrow indicates Yy1.

